# MicroMator: Open and Flexible Software for Reactive Microscopy

**DOI:** 10.1101/2021.03.12.435206

**Authors:** Zachary R Fox, Steven Fletcher, Achille Fraisse, Chetan Aditya, Sebastián Sosa-Carrillo, Sébastien Gilles, François Bertaux, Jakob Ruess, Gregory Batt

## Abstract

Microscopy image analysis has recently made enormous progress both in terms of accuracy and speed thanks to machine learning methods. This greatly facilitates the online adaptation of microscopy experimental plans using real-time information of the observed systems and their environments. Here we report MicroMator, an open and flexible software for defining and driving reactive microscopy experiments, and present applications to single-cell control and single-cell recombination.

## Introduction

Software for microscopy automation are essential to support reproducible high-throughput microscopy experiments^1^. Samples can now be routinely imaged using complex spatial and temporal patterns. Yet, in the overwhelming majority of cases, executions of experiments are still cast in stone at the beginning, with little to no possibility for human or computer-driven interventions during the experiments. This is all the more surprising given that image analysis has recently made a giant leap in terms of accuracy and rapidity thanks to deep learning methods, thus opening the way for implementing elaborate protocols. Software empowering microscopy with real-time adaptation capabilities is needed to exploit the full potential of automated microscopes.

Several dedicated microscopy software solutions have been developed for applications requiring real-time analysis. This is notably the case for the efficient scanning of large and complex microscopy samples (*eg,* Refs^2–6^). For other important applications, such as real-time control of cellular processes (*eg,* Refs^7–13^), results are generally obtained using ad hoc software solutions. Very few generic tools have been developed so far to facilitate the realization of complex, reactive microscopy experiments. One notable exception is Pycro-Manager^14^. This powerful framework is built on top of μManager, a widely-used software^15,16^ controlling a large range of microscopy hardware. In Pycro-Manager, reactive protocols are built from the ground up. Whereas this gives maximal flexibility, this also increases the difficulty to rapidly design experiments, especially for non-expert users. Moreover, no in-depth case studies demonstrating its practical applicability -and showing possible limitations-have been reported so far. One can also mention Cheetah, a simple to use Python library to support the development of real-time cybergenetic control platforms that combines microscopy imaging and microfluidics control^17^. In its current state, the possibilities to programmatically control the microscope appear limited.

## MicroMator Software

In this paper, we present MicroMator, a simple software solution supporting reactive microscopy experiments, and demonstrate its potential via two challenging case studies. In MicroMator, events play a fundamental role (Fig 1A). They consist of Triggers and Effects. They can be defined by the user in a flexible manner. Examples of triggers include “at the 10th frame”, “if more than 100 cells are in the field of view”, “if the fluorescence of the 3rd newborn cell exceeds 100 a.u.” and “if message !update position=10 frame=last is received from Discord”. Examples of effects include changing a microscope configuration, sending light in the field of view with a given pattern, actuating a microfluidic pump, or starting an optimization routine. Microscopy experiments are defined by a main image acquisition loop, that serves as a backbone for the experiment, and by event creation functions (Fig 1B). Naturally, the main acquisition loop itself can be modified by event effects in the course of the experiment. MicroMator is written in Python 3, is open-source and has a modular design. For controlling hardware, MicroMator primarily uses the powerful Python API of μManager pymmcore, but can also use other dedicated Python or web-based APIs provided by vendors, as done for our CellAsic ONIX microfluidic platform. Various types of analysis can be performed using dedicated software modules, such as on-line image analysis or real-time control and optimization. Communication modules can be used to interface MicroMator with digital distribution platforms such as Discord to track experiment progress and potential issues. Lastly, MicroMator leverages Python’s multiprocessing module to perform computations concurrently and possesses an extensive and customizable logging system, gathering logs of all modules in a unique file and fostering reproducible research (see SI Text).

**Figure 1.**
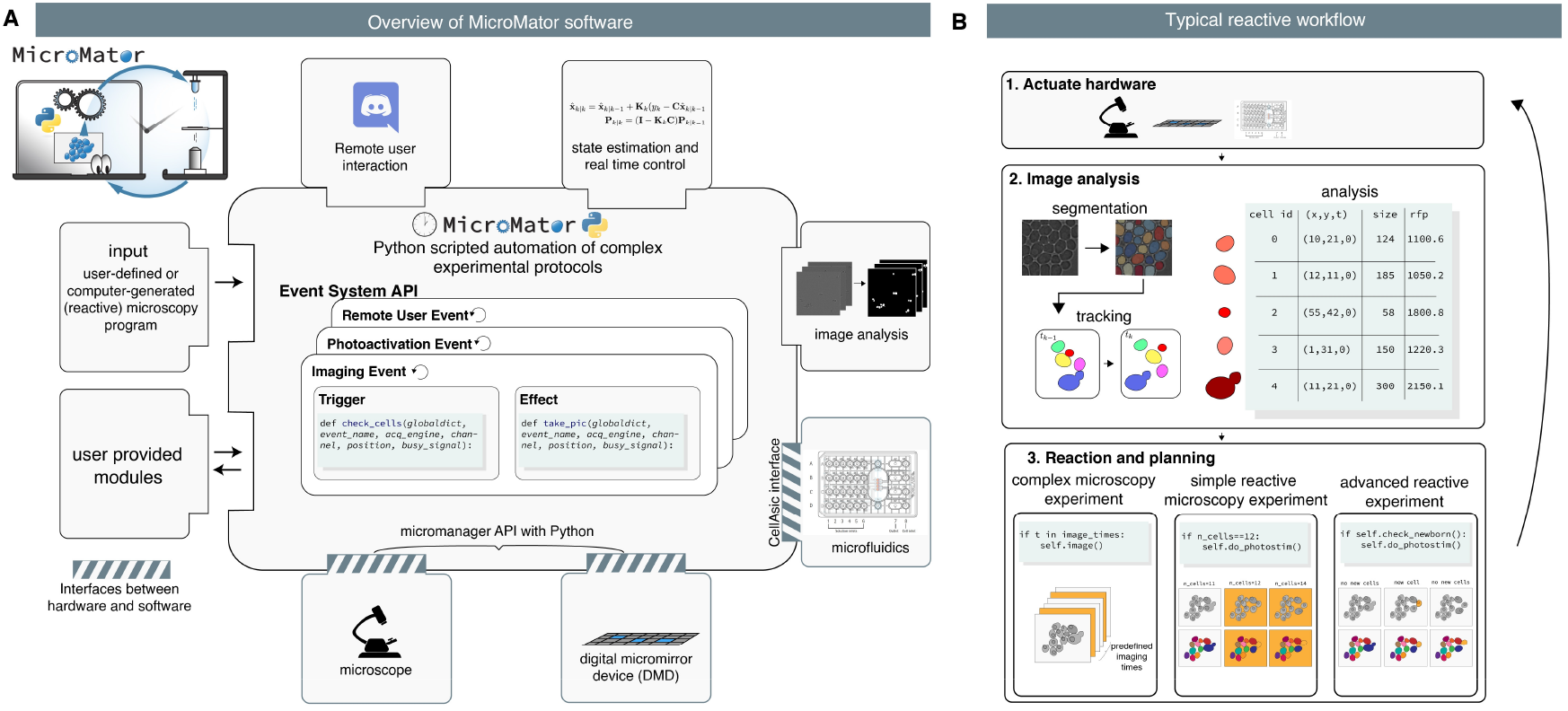
MicroMator overview. **A.** Modular software architecture. MicroMator consists of an extensible set of modules that control various hardware and software aspects of microscopy experiments and of a central core that handles user-defined events. It is written in the high-level programming language Python. **B.** Event-based reactive microscopy workflow. Imaging can be followed by online analysis of the samples. This typically involves segmentation, tracking, quantification of cell properties, and possibly advanced additional computations. Effects may then be triggered based on the result of the analysis. These may include the physical actuation of the hardware or the initiation of communications or of additional computations.

We also provide SegMator, a software which uses U-Net for bright-field yeast cell segmentation (provided by DeLTA^18^), and tracking using TrackPy^19^. U-Net is a convolutional neural network with a structure that excels at image segmentation^20^. U-Net can analyze dense fields of cells in a few seconds and with good accuracy (see SI Text and Fig S1 and S2 and Movie S1).

To showcase the full potential of reactive experiments performed with MicroMator, we designed experiments in which cellular processes are controlled in real-time. Single-cell stimulations are computed on line based on the cell state and/or position, demonstrating that reactive loops can be implemented at the level of individual cells. These experiments are inspired by previously-published studies^10,11,13,21^ and show how these could be repeated and further extended using generic software. We also provide a tutorial application in which cells with fluorescent proteins are imaged with increasing durations such that the measured intensity reaches a given threshold (Supplementary Text 1). This could typically be used to guarantee a good signal to noise ratio irrespectively of the initial fluorescence of the cells.

## Model predictive control of gene expression at the single-cell level in yeast

For our first application, we use the EL222 optogenetic system and the mScarletI fluorescent reporter to engineer light-responsive yeast cells (Fig 2A). Using real-time imaging, segmentation and cell tracking, different cells can be stimulated differently in the field of view using a digital micromirror device (DMD). Our goal is to implement different model predictive control (MPC) strategies for controlling the expression levels of a protein in a cell population. The cellular response of our engineered cells was characterized for different light stimulation profiles in the same experiment (Fig 2B). Only the most central part of the cell receives significant light stimulation. This erosion of the stimulation region helps improving the precision of single-cell light stimulations in dense cell regions because of illumination bleed-through of DMD systems (see SI Text and Fig S4). We then developed and calibrated an “average cell” (deterministic) and a “single cell” (stochastic) model of light-driven gene expression (SI Text and Fig S5, and S6).

**Figure 2.**
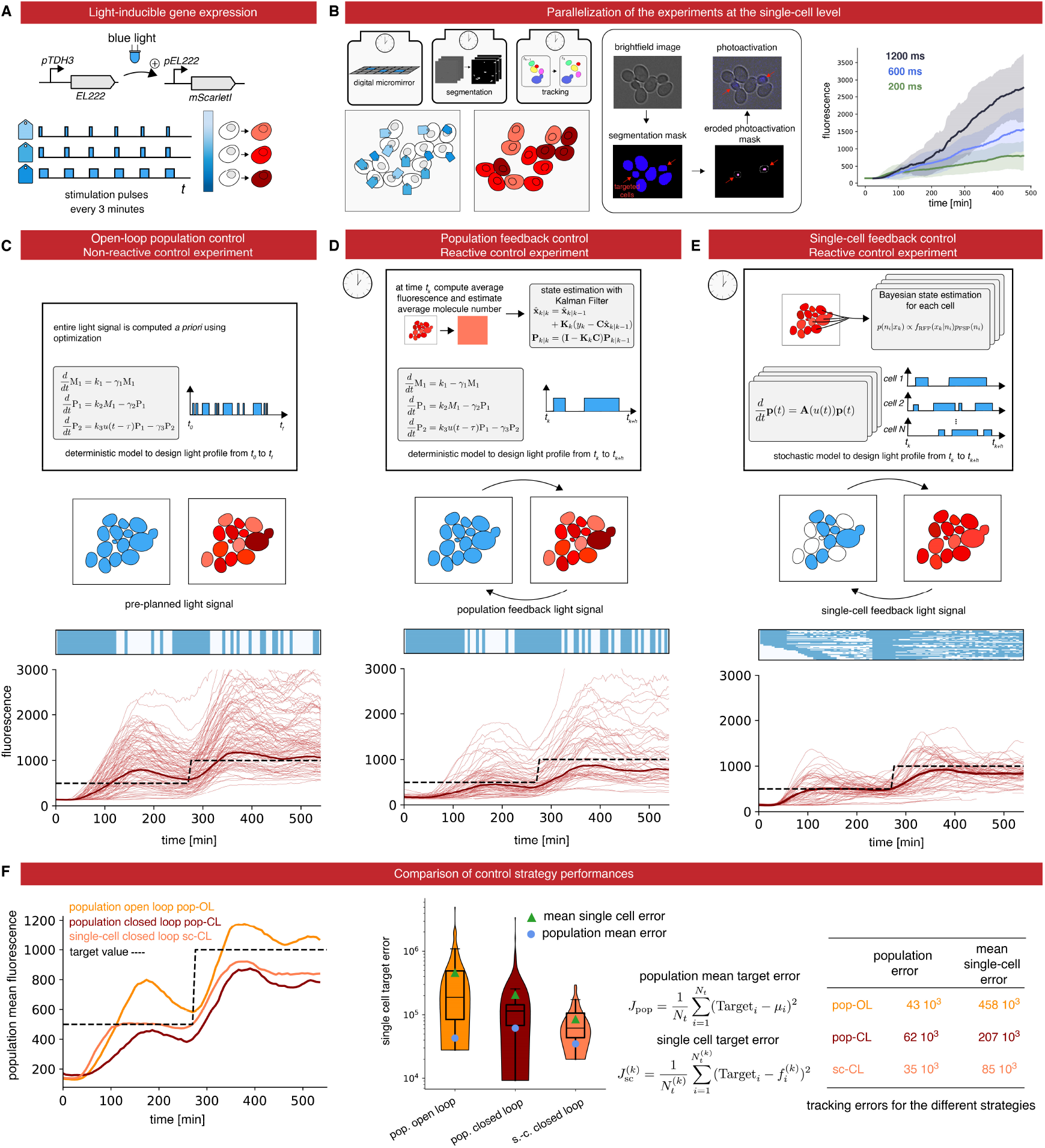
Control gene expression at the single cell level in yeast. **A.** The red fluorescent protein mScarletI is placed under the control of the light-responsive transcription factor EL222. **B**. To efficiently characterize cell responses to light stimulations, cells in the field of view are partitioned in 3 groups, each group being stimulated with a different temporal profile. Bright-field images are segmented and cells are tracked. Then, based on their groups, cells are stimulated during the appropriate time with eroded masks. The temporal evolution of the mean mScarletI fluorescence of the cells in the three groups is shown with envelopes indicating one standard deviation. **C.** Open-loop control experiment in which a model of the response of the cell population is used to precompute a light stimulation profile that drives the cell population to the target behavior. The application of the light profile leads to significant deviations from the target of the individual cell trajectories. **D.** Closed-loop control experiment in which the same model is used jointly with real-time observations of the population state to decide which light profile to apply to all cells, using a receding horizon strategy. **E.** A stochastic model of individual cell response is used jointly with single-cell observations to decide which light profile to apply to each cell. **F.** The different strategies have similar performances to drive the mean fluorescence to its target, but the single-cell feedback strategy is significantly better to drive individual cells to their target profiles.

In open loop control, the average cell model is used to precompute a temporal pattern of light stimulation so that cells follow a target behavior. This light pattern is then applied to all cells in the field of view (Fig 2C and Movie S2). In closed loop population-based control, the average cell model and the average of the measured fluorescence of cells are used by classical state estimators and model predictive controllers to compute in real-time the appropriate light stimulation to drive the mean fluorescence to its target (Fig 2D and Movie S3). Finally, in closed loop single-cell control, a stochastic model of gene expression and single-cell fluorescence measurements are used by advanced state estimators and controllers to compute in real-time the appropriate light stimulations to drive the fluorescence of each and every cell in the field of view to its target (Fig 2E and Movie S4). This control problem is quite challenging and needs to be solved for hundreds of cells in parallel. Advanced methods for numerical simulation and state estimation were essential (see SI Text and Fig S7).

Defining control performance as the time averaged deviation to target, we found that the single-cell control method leads to a modest reduction of error of the population averaged fluorescence but to a drastic improvement of the average error of the single-cell fluorescence (Fig 2F).

## Patterns of recombined yeast cells

For our second application, we constructed a light-driven artificial recombination system in yeast and employed different light stimulation strategies to obtain various structures of recombined cells.

We again used the EL222 optogenetic induction system but this time to drive the expression of the Cre recombinase. The Cre recombinase induces the expression of a fluorescent reporter, mCerulean, via an amplification step using the ATAF1 transcription factor (Fig 3A & 3B). This strain has been designed as in Ref ^22^.

**Figure 3.**
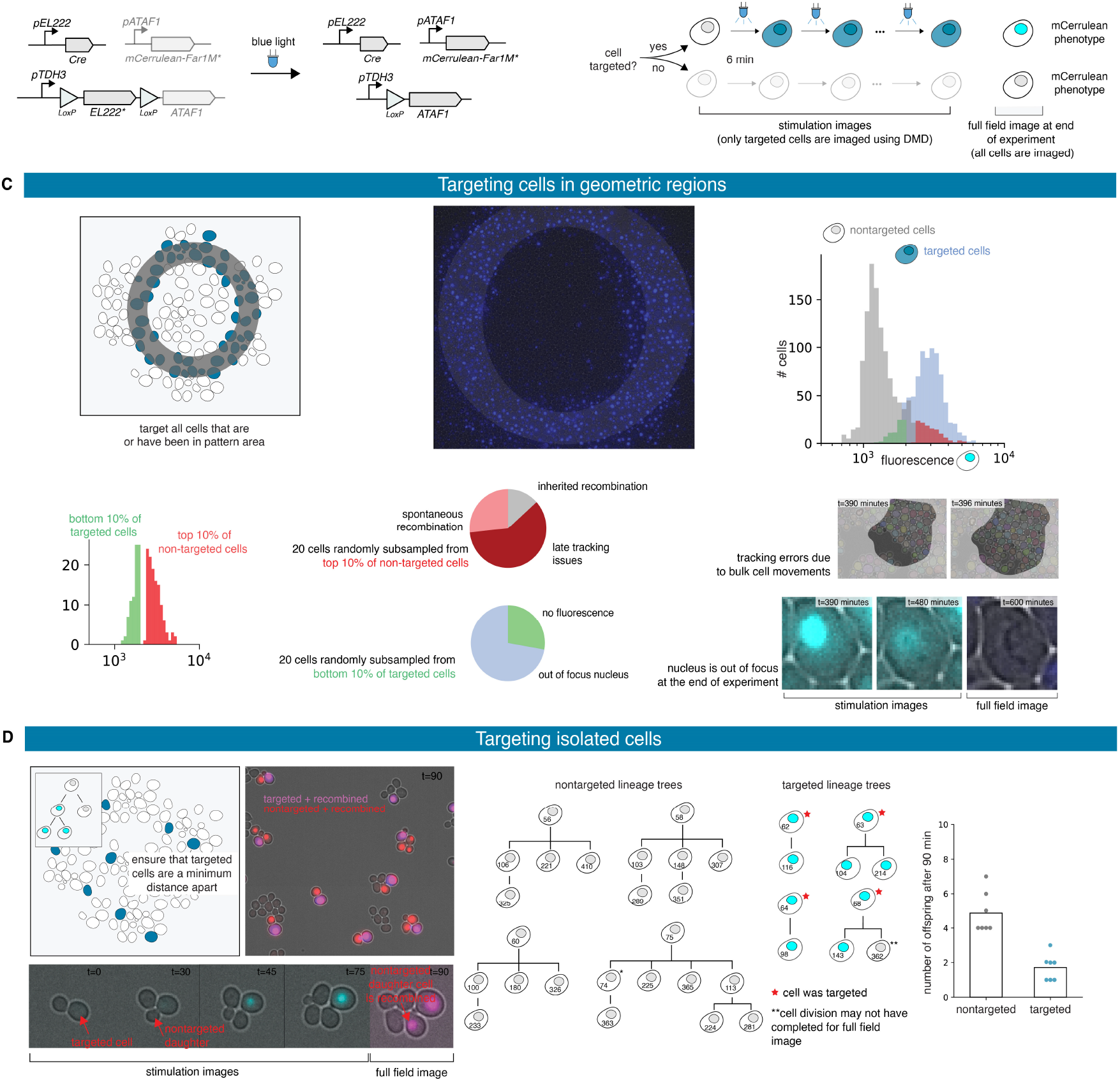
Patterns of recombined yeast cells. **A.** Upon light exposure, the Cre recombinase is expressed and triggers recombination, leading to the expression of ATAF1 and then of mCerulean-Far1M. Stars indicate nuclear localization of the protein. **B.** Targeted cells are stimulated for 1 second every 6 minutes until the end of the experiment. Fluorescence levels emitted by targeted cells can be recorded. At the end of the experiment, all cells are imaged and a recombined or non-recombined phenotype is attributed. **C.** A ring-like region in the field of view is selected at the beginning of the experiment and all cells entering the designated region at some time point are targeted for recombination. The distributions of the fluorescence levels of the targeted and non-targeted cells can be computed at the end of the experiment. The vast majority of cells present the expected phenotype and outliers can be further analyzed. **D.** Cells are dynamically selected such that no target cells are close to each other. Cell lineages of targeted and non-targeted cells can be manually reconstructed and statistics can be extracted.

Firstly, we applied a ring-like recombination signal. More specifically, every cell that was in the designated zone at any moment throughout the experiment has been targeted for recombination (Movie S5). As a result, we did obtain a ring-like pattern of recombined cells (Fig 3C). Experimental and biological limitations can be revealed by the analysis of the tails of the distributions of the recombination readout *(i.e.,* mCerulean fluorescence) within the cell populations (Fig 3C). For example, we found that some cells have been erroneously targeted for recombination because of tracking issues, and that only a few cells have not shown the recombined phenotype at the end of the experiment despite having being effectively targeted for recombination (Fig 3C and S9).

Secondly, we tried to create islets of recombined cells. To this end, we dynamically searched for cells that were far from any previously-targeted cell, and targeted these cells for recombination. To maximize the chances that the chosen cells do recombine, we tracked each chosen cell and targeted it repeatedly with light stimulations (Movie S4). Our strategy was effective in creating isolated microcolonies of recombined cells (Fig 3D). Analysis of the lineage trees of targeted cells and non-targeted cells confirmed that recombined cells have a slow growth phenotype. Previous works demonstrating optogenetically-driven recombination use static masks for light targeting^21^. Obtaining single-cell resolution as demonstrated in Fig 3D necessitates real-time image analysis and the use of reactive software.

## Discussion

We presented MicroMator together with two challenging applications demonstrating how this software can help using automated microscopy platforms to their full potential. Each application goes beyond the state of the art. We showed the first demonstration of control of protein expression at the single cell level in dense field of cells. This requires to jointly solve two challenges, namely obtaining sufficiently precise single-cell stimulations with DMDs and segmenting and tracking cells with sufficient accuracy over extended durations. We also provide the first demonstration of cell recombination targeted at the single-cell level, enabling single-cell resolution patterns. In comparison with Pycro-Manager, MicroMator uses the Micro-Manager GUI to create a main acquisition backbone for the experiment and reactive events are then used to enhance or even dynamically modify this initial plan. Events are created by default as separated threads and an extended logging system gathers messages from all modules that might be running in parallel. This structure provides robustness to real-time issues and facilitates error identification, two critical aspects for developing long and complex experiments.

Yet, we foresee that reactiveness in microscopy will primarily be used to enhance and automate classical experiments. Examples of simple use cases abound: triggering autofocus only when needed, dynamically adjusting the imaging condition to the signal strength, identifying novel regions of interest, or following the course of experiments via easily accessible online services (e.g. warning messages sent on Discord), to provide but a few examples. Thanks to its modular nature and to its use of a simple but powerful event system, MicroMator capacities can be conveniently expanded to drive novel hardware or perform a wide range of analyses. MicroMator is a relatively simple software extension that significantly empowers laboratory equipment that is present in most quantitative biology laboratory worldwide.

## Methods

### Software and data availability

MicroMator is an open-source software. It contains a core part and an extensible list of modules. The MicroMator core manages the user-specified events and also the metadata and logging system. The current list of modules includes a Microscope Controller module, an Image Analysis module, a Model Predictive Control module, and a Discord Bot module. The Microscope Controller module is an interface with the Python wrapper for MicroManager pymmcore. The Image Analysis module uses deep learning methods to segment yeast cells from bright-field images. It also uses an efficient algorithm for cell tracking. This module is also available as a standalone tool called SegMator. The Model Predictive Control module implements state estimation and model predictive control routines for deterministic and stochastic systems, at either the population or single-cell level. The Discord Bot module uses a web app running on the microscope’s computer and connected to the Discord communication system.

MicroMator, SegMator, event definitions for representative experiments (Fig 2E and S4), and data analysis code for the experiments (Fig 2E, 3D, and S4), as well as a tutorial example (Supplementary Text 1), can be found online: https://gitlab.inria.fr/InBio/Public/micromator. Raw and processed data for Fig 2C-E, 3C-D, and S4 are freely available on the zenodo repository: https://doi.org/10.5281/zenodo.4616659 (45GB).

Supplementary movies

- **Movie S1**: real-time_segmentation_and_tracking_with_SegMator.mov. Time-lapse movie showing the real-time segmentation and tracking quality obtained with SegMator. Left: bright-field image. Right: bright-field image overlaid with segmentation mask in cyan.
- **Movie S2**: optogenetic_control_of_gene_expression-Open_loop.mov. Time-lapse movie showing the response of the cells (mScarletI fluorescence) in an open-loop control experiment. Corresponds to Fig. 2C.
- **Movie S3**: optogenetic_control_of_gene_expression-Population_closed_loop.mov. Time-lapse movie showing the response of the cells (mScarletI fluorescence) in a population closed-loop control experiment. Corresponds to Fig. 2D.
- **Movie S4**: optogenetic_control_of_gene_expression-Single_cell_closed_loop.mov. Time-lapse movie showing the response of the cells (mScarletI fluorescence) in a single-cell closed-loop control experiment. Corresponds to Fig. 2E.
- **Movie S5**: single_cell_recombination-Islets.mov. Time-lapse movie showing the light signal sent to cells in order to create small islets of recombined cells. Corresponds to Fig 3C.
- **Movie S6**: single_cell_recombination-Ring.mov. (Left) Time-lapse movie showing the light signal sent to cells in order to recombine all cells that have been at one moment in a ring-like pattern. (Right) Image showing the recombined state of the cells at the end of the experiment. Corresponds to Fig 3D.

### Genetic constructions and yeast strains

All plasmids and strains were constructed using the *Yeast Tool Kit,* a modular cloning framework for yeast synthetic biology^23^, the common laboratory strain BY4741 (Euroscarf), and the EL222 optogenetic system^24^. The light responsive strain (IB44) harbors a constitutively expressed EL222 light-responsive transcription factor (NLS-VP16AD-EL222) and an EL222-responsive promoter (5xBS-CYC180pr) driving the expression of the mScarletI protein. The IB44 strain genotype is MATa his3Δ1 leu2Δ0::5xBS-CYC180pr-mScarletI-Leu2 met15Δ0 ura3Δ::NLS-VP16AD-EL222-URA3. The recombining strain (IB237) harbors a constitutively expressed EL222 light-responsive transcription factor (NLS-VP16AD-EL222) floxed between two LoxP sites that upon recombination expresses the ATAF1 transcription factor. This factor expresses (pATAF1_4x) in turn the mCerulean fluorescent protein fused to a constitutively active Far1 protein (FAR1M_mCerulean). Lastly, the strain also harbors the Cre recombinase placed under the control of an EL222-responsive promoter (5BS-Gal1pr). The IB237 strain genotype is MATa his3Δ1::pATAF1_4x-FAR1M_mCerulean-tDIT1-HIS3 leu2Δ::5BS-Gal1pr-CRE-tENO1-LEU2 met15Δ0 ura3Δ:: pTDH3-LoxP-NLS-VP16AD-EL222-tENO1-LoxP-ATAF1-tTDH1-URA3. Lastly, we also used the IB84 strain as a constitutive 3-color strain to characterize DMD precision. The genotype of this strain is MATa his3Δ1 leu2Δ0::pTDH3-mCerulean-tTDH1-pTDH3-NeonGreen-tTDH1-pTDH3-mScarlet-tTDH1-LEU2 met15Δ0 ura3Δ:: NLS-VP16AD-EL222-URA3.

### Culture preparation

Cells were grown at 30°C in synthetic medium (SD) consisting of 2% glucose, low fluorescence yeast nitrogen base (Formedium CYN6510), and complete supplement mixture of amino acids and nucleotides (Formedium DCS0019). For each experiment, cells were grown overnight in SC media at 30°C, then diluted 50 times and grown for 4 to 5 hours before being loaded in microfluidic plates.

### Microscopy setup, microfluidics and imaging

Images were taken under a Leica DMi8 inverted microscope (Leica Microsystems) with a ×63 oil-immersion objective (HC PL APO), an LTM200 V3 scanning stage, and an sCMOS camera Zyla 4.2 (ANDOR). Bright-field images were acquired using a 12V LED light source from Leica Microsystems. Fluorescence images were acquired using a pE-4000 light source from CoolLED and the following filter cubes: EX:436/20nm DM:455nm EM:480/40nm (CFP), EX:500/20nm DM:515nm EM:535/30nm (YFP), and EX:546/10nm DM:560nm EM:585/40nm (RHOD) from Leica Microsystems. Light stimulation was performed using the pE-4000 light source and the CFP filter. Spatially-resolved illuminations were obtained thanks to a digital mirror device (DMD) reflecting the light of a pE-4000 light source. We used a MOSAIC3 DMD from ANDOR. The device is used both for targeted fluorescence imaging and for optogenetic stimulations. A CellASIC ONIX2 system (Merck) was used together with the Y04C CellASIC microfluidic plates to grow yeast cells in monolayers. Media flow was maintained by a 7.5 kPa pressure gradient. The media was the same as for pre-culture. The temperature was maintained at 30°C by an opaque environmental box and a temperature controller 2000-2, both from PECON. The microscope was operated using MicroMator.

### Model predictive control of gene expression

To compare single-cell and population control strategies, we developed stochastic and deterministic models of gene expression. Both have been calibrated with respect to the dataset presented in Fig 2B and Fig S5. For population control, we used the deterministic model, assumed Gaussian measurement noise and used a Kalman filter for state estimation. Each model assumes a deterministic delay between the time the light signal is applied and the time protein production is effective. For MPC, fluorescence measurements were taken every 6 minutes and we considered receding time horizons of 24 minutes. The controller explores the set of light stimulation profiles in which a 1000ms light stimulation is either applied or not for each measurement time interval, and selects the profile minimizing mean square deviations. For tracking purposes, brightfield measurements were taken every 3 minutes. For single-cell control, we used the stochastic model and simulated the cell behavior using a finite state projection approximation. For each and every cell, state estimation is performed using a Bayesian approach which conditions the probability distribution for each cell on the most recent measurement, and light stimulation profiles are selected using the approach outlined above and the expected absolute deviation as selection criterion. More information is provided in SI Text. Box plots of Figure 2F indicate the lower quartile, the median, and the upper quartile of the data, with the whiskers corresponding to 1.5 interquartile ranges.

## Acknowledgments

This work was supported by ANR grants CyberCircuits (ANR-18-CE91-0002), MEMIP (ANR-16-CE33-0018), and Cogex (ANR-16-CE12-0025), by the H2020 Fet-Open COSY-BIO grant (grant agreement no. 766840) and by the Inria IPL grant COSY. We acknowledge the support of the U.S. Department of Energy through the LANL/LDRD Program and the Center for Non Linear Studies for this work.

## Author contributions

S.F. developed the MicroMator software and performed experiments. Z.F. wrote the image analysis and real-time control modules, and analyzed the data. A.F. performed experiments and helped develop the MicroMator software and analyze the data. C.A. developed strains and helped perform experiments. S.S.-C. developed strains. S.G. helped with software development. F.B. helped with software development and platform integration. J.R. helped with controller development. Z.F., F.B., J.R., and G.B. supervised the project. Z.F. and G.B. wrote the manuscript with input from all authors.

## References

1. Eisenstein, M. Smart solutions for automated imaging. Nat Methods 17, 1075–1079 (2020).

2. Conrad, C. et al. Micropilot: automation of fluorescence microscopy-based imaging for systems biology. Nat Methods 8, 246–249 (2011).

3. Carro, A., Perez-Martinez, M., Soriano, J., Pisano, D. G. & Megias, D. iMSRC: converting a standard automated microscope into an intelligent screening platform. Sci Rep 5, 10502 (2015).

4. Pinkard, H., Stuurman, N., Corbin, K., Vale, R. & Krummel, M. F. Micro-Magellan: open-source, sample-adaptive, acquisition software for optical microscopy. Nat Methods 13, 807–809 (2016).

5. Li, T. et al. MAARS: a novel high-content acquisition software for the analysis of mitotic defects in fission yeast. MBoC 28, 1601–1611 (2017).

6. Politi, A. Z. et al. Quantitative mapping of fluorescently tagged cellular proteins using FCS-calibrated four-dimensional imaging. Nat Protoc 13, 1445–1464 (2018).

7. Toettcher, J. E., Gong, D., Lim, W. A. & Weiner, O. D. Light-based feedback for controlling intracellular signaling dynamics. Nat Methods 8, 837–839 (2011).

8. Uhlendorf, J. et al. Long-term model predictive control of gene expression at the population and single-cell levels. PNAS 109, 14271–14276 (2012).

9. Lugagne, J.-B. et al. Balancing a genetic toggle switch by real-time feedback control and periodic forcing. Nat Commun 8, 1671 (2017).

10. Chait, R., Ruess, J., Bergmiller, T., Tkačik, G. & Guet, C. C. Shaping bacterial population behavior through computer-interfaced control of individual cells. Nat Commun 8, 1535 (2017).

11. Rullan, M., Benzinger, D., Schmidt, G. W., Milias-Argeitis, A. & Khammash, M. An Optogenetic Platform for Real-Time, Single-Cell Interrogation of Stochastic Transcriptional Regulation. Molecular Cell 70, 745–756.e6 (2018).

12. Perrino, G., Wilson, C., Santorelli, M. & di Bernardo, D. Quantitative Characterization of α-Synuclein Aggregation in Living Cells through Automated Microfluidics Feedback Control. Cell Reports 27, 916–927.e5 (2019).

13. Perkins, M. L., Benzinger, D., Arcak, M. & Khammash, M. Cell-in-the-loop pattern formation with optogenetically emulated cell-to-cell signaling. Nat Commun 11, 1355 (2020).

14. Pinkard, H. et al. Pycro-Manager: open-source software for customized and reproducible microscope control. Nat Methods (2021) doi:10.1038/s41592-021-01087-6.

15. Edelstein, A., Amodaj, N., Hoover, K., Vale, R. & Stuurman, N. Computer Control of Microscopes Using μManager. Current Protocols in Molecular Biology 92, 14.20.1–14.20.17 (2010).

16. Edelstein, A. D. et al. Advanced methods of microscope control using μManager software. J Biol Methods 1, 10 (2014).

17. Pedone, E. et al. Cheetah: A Computational Toolkit for Cybergenetic Control. ACS Synth. Biol. acssynbio.0c00463 (2021) doi:10.1021/acssynbio.0c00463.

18. Lugagne, J.-B., Lin, H. & Dunlop, M. J. DeLTA: Automated cell segmentation, tracking, and lineage reconstruction using deep learning. PLoS Comput Biol 16, e1007673 (2020).

19. Allan, D. et al. soft-matter/trackpy: Trackpy v0.4.2. (Zenodo, 2019). doi:10.5281/ZENODO.3492186.

20. Falk, T. et al. U-Net: deep learning for cell counting, detection, and morphometry. Nat Methods 16, 67–70 (2019).

21. Polstein, L. R., Juhas, M., Hanna, G., Bursac, N. & Gersbach, C. A. An Engineered Optogenetic Switch for Spatiotemporal Control of Gene Expression, Cell Differentiation, and Tissue Morphogenesis. ACS Synth. Biol. 6, 2003–2013 (2017).

22. Aditya, C., Bertaux, F., Batt, G. & Ruess, J. A light tunable differentiation system for the creation and control of consortia in yeast. bioRxiv 2021.06.09.447744 (2021).

23. Lee, M. E., DeLoache, W. C., Cervantes, B. & Dueber, J. E. A Highly Characterized Yeast Toolkit for Modular, Multipart Assembly. ACS Synth Biol 12 (2015).

24. Benzinger, D. & Khammash, M. Pulsatile inputs achieve tunable attenuation of gene expression variability and graded multi-gene regulation. Nat Commun 9, 3521 (2018).

