## Supplementary material for "MicroMator: Open and Flexible Software for Reactive Microscopy"

#### Contents

**1 Software Description** **2**

**2 Example of a simple reactive experiment with MicroMator** **7**

**3 Analysis of DMD Precision Using Three Color Fluorescent Microscopy** **9**

**4 Model Predictive Control of Optogenetically Induced Protein Production** **9**

**5 Data Analysis** **18**

### 1 Software Description

The software was designed around different modules, such as image analysis, microscope control and a custom event system, which all come together and allows the user to design and create interactive feedback loops. The other core features of MicroMator's modular design include a clear and flexible API, a set of reactive image analysis modules, a microscope module that uses Micro-Manager's generic and open source framework and its Python library, pymmcore, and an extensive logging system and metadata management system.

MicroMator is open-source, and can be adapted to the user's needs. The main interface point is the event system, which is described in more detail below. One may also design their own modules, where users can do more complex tasks (see our model predictive control module `Stoched` as an example), but a more complete understanding of how MicroMator works is required to integrate these types of modules. Here we present brief descriptions of the software itself, MicroMator Core, along with the modules we have written to perform the experiments in the paper. The software is available on Gitlab, which includes the modules required to replicate the experiments presented here and in the main text. More information about using the software is available in the README.md file on the Gitlab page.

#### MicroMator Core

MicroMator Core is the system that links together different modules in a unified framework. Importantly, it manages the events of the system, which are broken into Triggers and Actions. Events are constantly checked by MicroMator Core. Users can write their own events that then interface with the rest of the system. A Trigger is a Python class that contains a method named `check`, which returns a Boolean indicating if a given action should take place. Each event is called throughout the experiment, and at that time the `trigger.check()` is called. If the check returns `True`, then the corresponding Action is called. Actions should have a method `act`, which describes what should happen when the event has been triggered. We use multithreading to constantly check if each event is available (i.e. it is not currently doing something) or busy. If a check returns `True` while an event is busy, the event is queued.

Importantly, MicroMator Core also handles all of the logging of the system. It records what each module is doing, and saves all of the logs into a consistent format. The log files are critical to debugging the system. For example, in an experiment with multiple fields of view, each of which is performing a different light stimulation, the logging system will record what the software is doing at each position, which modules were used, and any module or event specific action. Next, we will describe the modules that are included in MicroMator.

#### Microscope Module

The Microscope Module is in charge of acquiring the raw images, according to the acquisition protocol and the different channels and positions the user has preset at the start of the experiment. Micro-Manager's GUI is used to generate the experiment's initial conditions, such as the number of time steps, the different channels and the interesting positions for the user. A custom implementation of Micro-Manager's pymmcore API will read, save this data and then execute the acquisition.

#### Image Analysis Module

A critical component of reactive microscopy is the notion of online image analysis. It allows the user to program experiments that adapt to the behavior of the cells in real time. MicroMator includes an Image analysis module, which can be called every frame, or at a subset of frames, similar to the events system described below. As part of developing MicroMator, we developed an Image Analysis module which we called `SegMator`. This module uses modern deep learning approaches to segment yeast cells based on brightfield images. It can then optionally track segmented cells across frames. `SegMator` generates and iteratively updates a Pandas dataframe as data is collected. Depending on the experiment, this data file contains identities for each cell, as well as the position, segmentation contour, cell size, fluorescence and other individual cell data that may be used to be reactive within the experiment. We next describe these segmentation and tracking aspects.

##### Cell Segmentation

Brightfield images were segmented using the U-Net, a convolutional neural network with a structure that excels at image segmentation [1, 2, 3]. An essential component of our system is to segment and track cells in real time within multiple fields of view. A key advantage of deep learning based approaches is that they are computationally expensive to train, but evaluations of a single 1024x1024 image are very fast and can accurately segment dense fields of cells in under a couple of seconds. We started with the pretrained segmentation network for yeast provided in the DeLTA package [2], which was implemented with the popular neural network packages Keras and Tensorflow [4, 5]. We developed our own preprocessing pipeline to implement the weight maps in Python, and created GUI's using the napari viewer [6] to create new masks for training. We used these tools to develop new training images for the network on our brightfield images. We found that relatively few (around 50) new training images were needed to adapt the network to our setup. This is likely because we started with a network which was already optimized for segmenting yeast, and the on-the-fly Python generators provided by DeLTA [2] which automatically apply random transformations to flip/shift/rotate the images and even simulate different illumination conditions. Code and our pretrained model files are available on Gitlab. Segmentation examples are shown in Figs. S1 and S2.

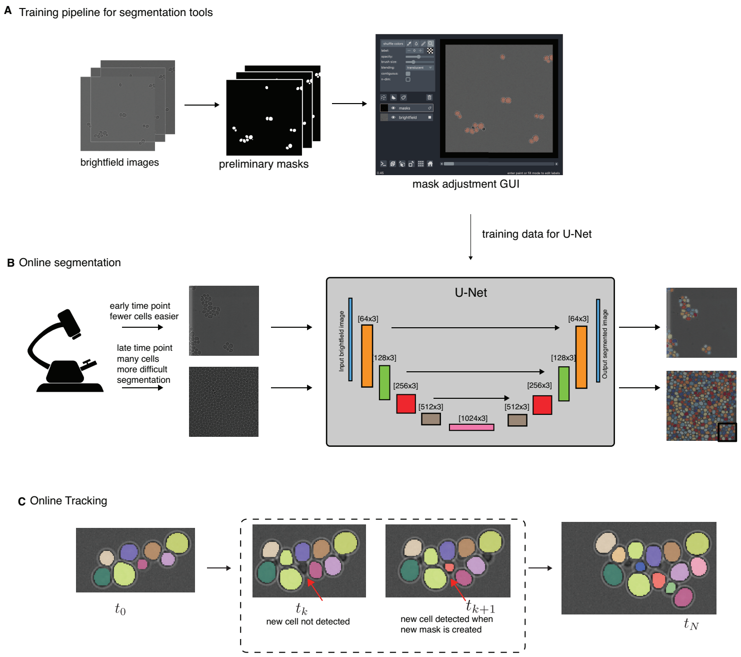


Figure S1: Overview of the segmentation and single cell tracking workflow. (A) Brightfield images were segemented using the previous iteration of U-Net, and then mask images were adjusted using a custom napari GUI. The network was then retrained on the new images and validated on new test images. (B) The online segmentation of cells in real time using U-Net. (C) Example of the tracking pipeline applied.

#### Cell Tracking

After masks were created using the neural network, they were linked using Trackpy software. We used the particle tracking algorithm from Crocker and Grier [7]. We verified the tracking algorithm quality by tracking the cells by eye from a movie to get an idea of how long cells are correctly tracked on average. We found that some cells "hop" long distances between measurement times (every 3 minutes). These cells sometimes disappear completely (i.e. they are washed away) or they reappear at a far distance. We do not expect to accurately track such cells, and therefore do not count them as errors in our tracking algorithm. We found that the vast majority of cells were correctly tracked over the entire time horizon while some cells were not, as summarized in Fig. S2. We also found that the tracking is affected by the time between frames. For all of the EL222/mScarlet characterization and control experiments, we had three minutes between frames, which was sufficient for accurate tracking, as shown in Fig. S1. However, for the recombination experiments we wished to limit the amount of light exposure, and we therefore took images every 6 minutes. While tracking

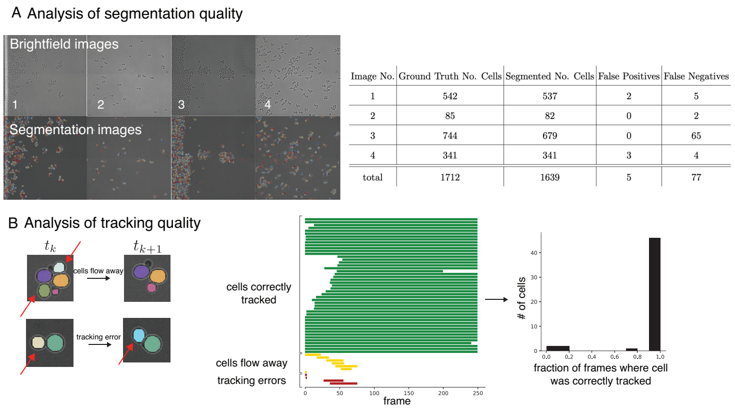


| Image No. | Ground Truth No. Cells | Segmented No. Cells | False Positives | False Negatives |
|---|---|---|---|---|
| 1 | 542 | 537 | 2 | 5 |
| 2 | 85 | 82 | 0 | 2 |
| 3 | 744 | 679 | 0 | 65 |
| 4 | 341 | 341 | 3 | 4 |
| total | 1712 | 1639 | 5 | 77 |

Figure S2: Validation of the segmentation and tracking pipeline. (A) Representative brightfield images and segmentation images. The table summarizes the segmentation quality for each sample image. (B) Validation of the tracking errors. Track lengths on a representative movie are shown in the middle panel. The fractions of the traces that correspond to correct trackings are represented in the rightmost panel.

was still possible when the field was sparse, for dense fields with bulk movement tracking quality was significantly diminished, as shown in Fig. S9.

#### Single-Cell Control Module

This is the module which implements our model predictive control for EL222 population and single-cell models, which we call `Stoched`. The module works by sorting the entire population at a given time into on or off cells according the control program, which is described below in Section 4. For the population control, by definition all cells are assigned as either on or off. However, for the single cell control experiments, (see Fig. 2 in the main text) different individual cells are assigned identities, and tracking is required. The module interfaces with individual cell identities using a main data file, which is a Pandas dataframe that is iteratively updated with tracking information, segmentation information, and fluorescence measurements. As mentioned above, this file is generated and updated in real time using the `SegMator` module. This file is the input to the control algorithms for both single-cell and population based control. The module then interfaces with the DMD to apply light to selected cells.

For the individual cell control, each cell is running its own stochastic model. Because we took a computationally expensive approach to solving the model using the Finite State Projection (FSP) (see Section 4 for mathematical details), it would be computationally infeasible to solve the control problem for each cell in series. We developed `Stoched` so that when one has multiple controllers the control problem can be performed across multiple processes. We used Python's multiprocessing module to spawn a new process for each controller, which allowed us to solve the control problem within the 6 minutes we had between measurement updates for each cell.

#### Discord Bot Description

The discord bot is a web app running on the microscope's computer and connected to the discord communication software. It allows the user to check the state of the experiment from anywhere with an internet connection. Through chat commands the user can inspect the last images taken by MicroMator and have access to MicroMator's log files to see if an error occurred. Furthermore, the bot inspects the Python logs every frame looking for exceptions and errors, and will notify the user on discord if such errors are detected. Currently, the bot is capable of reading MicroMator data but not of modifying or cancelling the experiment as the experiment continues.

#### Metadata Management and Logging

MicroMator also feature some metadata management and stores all the parameters in a few separate objects:

- `Protocol` class stores all the information regarding the microscope acquisition loop.
- `Channel` class stores all the channels the user will need for an experiment.
- `Position` class stores all the different positions the user saves for the experiment.

It is from these objects that the various modules get their parameters. When the experiment starts, MicroMator creates various folders and subfolders to store all the relevant information that is generated during the experiment, i.e. raw images, analyzed images, protocols, positions, and logs.

One advantage of MicroMator is the centralized logging capabilities. All of the included modules extensively write to a single log-file. User created modules are also able to write to the same `logger` object.

#### Writing your own Modules

If the default modules are not suited for the strain or the setup, the user can create their own module using the MicroMator API. On the Gitlab page, we provide interfaces to CellStar [8] and to a simple `numpy` based image analyzer as examples of interfaces to other modules.

### 2 Example of a simple reactive experiment with MicroMator

We describe an experiment aiming at imaging objects of poorly-known fluorescence. The proposed strategy is to gradually increase exposure time until obtaining a strong enough fluorescence signal. Concretely, we start by exposing fluorescent cells during 10 ms and repeatedly increase the exposure time by 50 ms until the mean pixel fluorescence value reaches the (arbitrary) threshold of 1600.

This is implemented by having MicroMator using the Analysis module to compute the mean pixel fluorescence and creating events that increase the exposure duration if needed. All the needed files can be found on the MicroMator gitlab.

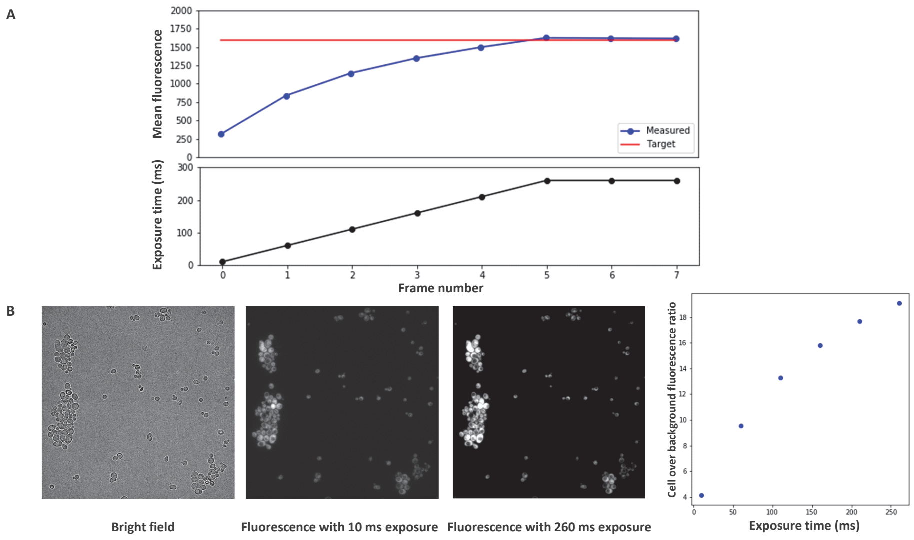


Figure S3: A simple reactive experiment with MicroMator. (A) After every time frame, the mean pixel fluorescence of the field of view is computed (blue), and if this value is below a given threshold (red, 1600 in this example), the exposure time is increased by by 50 ms (black, initial value is 10 ms). Frames are taken every 10 s. (B) Representative images of the cells taken in bright field and in fluorescence (YFP channel) for two different exposure times (frames 0 and 6). Yeast cells (*S. cerevisiae*) with constitutive mNeonGreen expression were imaged. One can check that the signal to noise ratio, defined here as the ratio between mean intracellular fluorescence and mean background fluorescence, increases with increasing exposure times.

The results of this experiment run by Micromator are presented in Fig. S3A. Representative images, together with a simple analysis of the signal to noise ratio, are presented in Fig. S3B. This simple example shows how one can ensure good imaging conditions for objects of a priori poorly-known fluorescence without bleaching the sample.

We will now present how to build an event object that will check the fluorescence, and take actions if needed.

##### The trigger

The trigger is fairly simple, we need to fetch the result of the numpy analysis and compare it to the threshold, returning True if the fluorescence is below and False otherwise.

```
class trigger_fluorescence_below_treshold:
    '''triggers if the fluorescence value measured by numpy is below the threshold
    '''
    def __init__(self, mmc, treshold):
        '''Args:
            mmc(object): the MMCore object
            threshold: a number (int or float)
        '''
        self.mmc = mmc
        self.treshold = treshold    # given as argument when creating the event

    def check(self, globaldict):    # check()  must take (self, globaldict) as args
       '''This check checks if the fluorescence value measured by numpy is below treshold
        '''
        numpy_data_path = os.path.join(self.mmc.folder_manager_obj.analysis_path,'data_0.csv')
        data = pd.read_csv(numpy_data_path)
        v = float(data['YFP-DIRECT'])
        print("TRIGGER: fluo value with current exposure: ", v, " wanted value: ",self.treshold)
        if v < self.treshold: #comparing last frame fluo value to our treshold
            return True
        else:
            return False
```

##### The effect

Here we need to modify the mmc.protocol object to raise the exposure by the chosen value. Note that this will change the parameters of the main acquisition loop.

```
class effect_augment_Exposure:
    '''this class augments the exposure of channel 1
    '''
    def __init__(self, mmc, value):
        '''Args:
            mmc(object): the MMCore object
            value(float): how much we raise the exposure
        '''
        self.mmc = mmc
        self.value = value

    def act(self, globaldict):
        '''this effector changes the exposure'''
        self.mmc.protocol.channel_list[1].exposure = self.mmc.protocol.channel_list[1].expo
sure + self.value    #editing the mmc.protocol object to change the acquisition parameters
```

##### The event creation function

The event creator file must have this function that returns a list of events. The events are built using the event class and the trigger and effect described earlier. This function will be launched at each time frame, and will effectively create the event.

```python
def create_events(mmc, globaldict):
    '''this is the functions that users will use to define their events
    Args:
        mmc(object): the MMCorePy object
        globaldict(dict): the dictionnary with all the global signals
    '''
    #list where all the created events are put
    events_list = []
    mytrigger = trigger_fluorescence_below_treshold(mmc, 1600)
    myeffect = effect_augment_Exposure(mmc, 50)

    event = manager.Event(mytrigger, myeffect, globaldict, name='exposure adjustment')
    events_list.append(event)
    return events_list
```

### 3 Analysis of DMD Precision Using Three Color Fluorescent Microscopy

To either image or activate single cells using the DMD, it is critical to understand the precision of the device. To do so, we targeted for illumination a subset of the cells and quantified the fluorescence in the targeted and in the non-targeted cells. We also tested the effect on bleedthrough of eroding the single-cell illumination masks. More specifically, we used a yeast strain with mCerulean, mNeonGreen, and mScarletI constitutively expressed (Fig. S4A). Then at each measurement time, we select three groups of cells, each to be imaged in a given color. Each cell is "tagged" in one color (Fig. S4B). Because all cells are expressing the three proteins, any accidental activation or bleedthrough in one imaging channel will show up as an off-target cell becoming illuminated. We measured the fluorescence values in all cells in each of the three channels using masks that were eroded to various degrees. A 75% erosion of the mask means that 25% of the area of the cell was retained for illumination. Sample microscopy images are shown in Fig. S4D. We measured the fluorescence in the target cells and in the off-target cells for 75%, 66%, 50%, 33%, and 25% erosion levels. The measured single-cell fluorescence values correspond to mean cell pixel intensities. From these experiments we determined that 33% erosion was sufficient to limit bleedthrough, which is critical when we photostimulate individual cells (Fig. S4E).

### 4 Model Predictive Control of Optogenetically Induced Protein Production

#### Characterization of Single-Cell EL222 Activation

Because we want to use the optogenetic system to control fluorescence values in individual cells, we performed a calibration experiment in which we randomly assigned cells in the field of view to receive 0, 200, 500, or 2000 ms of photostimulation. Cell masks were eroded to 33% of their original area to limit bleedthrough. When new cells were born, they were assigned a stimulation using our cell-sorter event, such that the population of cells in each bin was approximately even. Cells were stimulated every three minutes. We used these

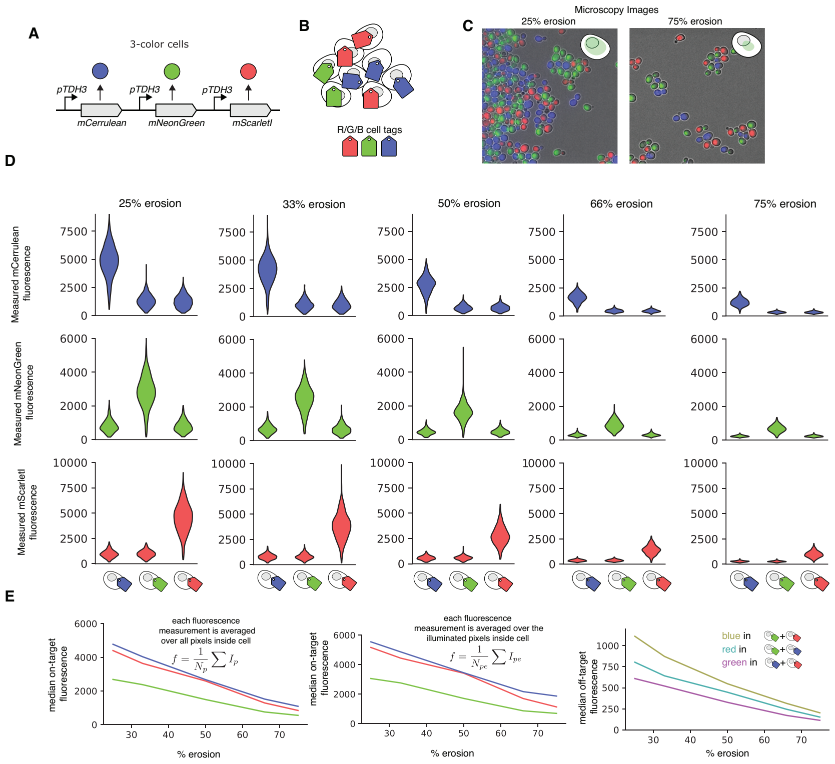

Figure S4: Effect of erosion on DMD precision imaging strategies. (A) Engineered *S. cerevisae* with three constitutively expressed fluorescent proteins. (B) Different individual cells are randomly assigned different color tags. (C) Tagged cells are illuminated in their channel using an eroded mask, and the fluorescence of all cells is recorded. The presented images are the merge of 3 single-color images and a brightfield image. (D) Fluorescence of all cells, targeted or not, in all fluorescence channels and for 5 different levels of erosion of the illumination mask. (E) Impact of erosion on on-target and off-target cell fluorescence. The median fluorescence of the targeted cells, the median fluorescence of the targeted areas in cells, and the median fluorescence of the non-targeted cells are represented as a function of the erosion of the illumination mask from left to right.

experiments as a proof of concept-to show that we could differentially stimulate the production of protein in individual cells.

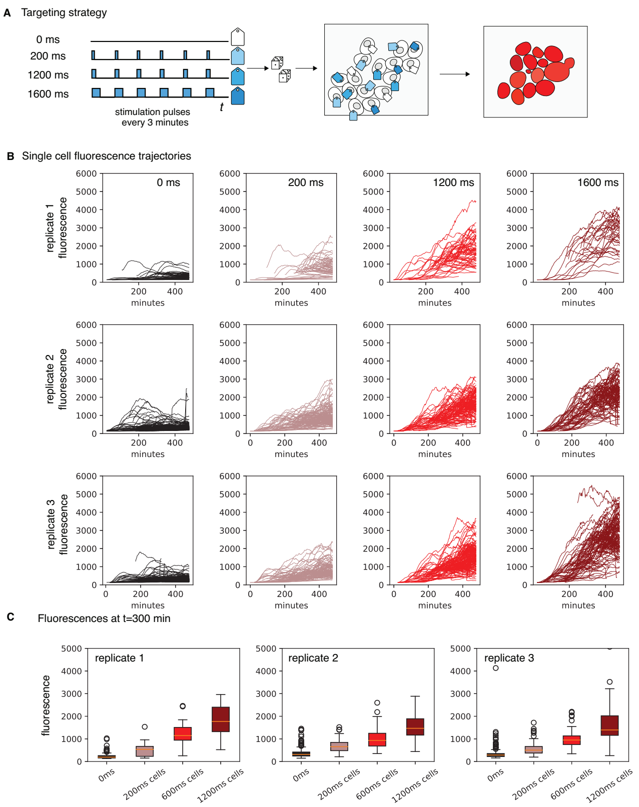


Figure S5: Single-cell photostimulation characterized in a single field of view. (A) Graphical description of the single-cell targeting for the system. (B) Time series of individual cell fluorescences in each photostimulation group across replicates. (C) Box plots showing the single-cell fluorescences at 300 minutes.

#### Stochastic Model of Protein Expression

We work specifically with the EL222 optogenetic system. In this system, EL222 proteins are created and degraded/diluted at all times within the cell because of stochastic protein production and degradation and cell growth. Under blue light photostimulation, EL222 molecules dimerize and then become activated transcription factors, which bind to the pEL222 promoter, activating protein production. We model the system of light-induced protein production using the following set of biochemical reactions, describing the constitutive transcription and translation of EL222 molecules, which we call $X$, and the production of fluorescent protein, $Y$, together with their degradation/dilution,

$$\begin{aligned}
&\varnothing \xrightarrow{k_1} B \times X \\
&X \xrightarrow{\gamma_1} \varnothing \\
&\varnothing \xrightarrow{k_2 u(t-\tau)} Y \\
&Y \xrightarrow{\gamma_2} \varnothing.
\end{aligned} \tag{1}$$

$B$ is a random variable which describes a "burst" of EL222 expression, and is geometrically distributed [9]. This model is valid when mRNA are short-lived compared to protein, and allows us to simplify the model. The control input, $u(t)$, is Boolean and represents the light being either on or off. We assume a delay $\tau$ between the time the light is applied and fluorescent protein is produced. The propensities of the system are:

$$\begin{aligned}
w_1(x_i) &= k_1(1-b)^{x_i-1}b, & w_2(x_i) &= \gamma_1 x_i \\
w_3(y_i) &= k_2 x_i u(t-\tau), & w_4(y_i) &= \gamma_3 y_i.
\end{aligned}$$

The mean burst size of the geometric distribution is given by $1/b$. The infinitesimal generator can therefore be written

$$\mathbf{A}_{i,j} = \begin{cases} -\sum_{k=1}^{4} w_k(x_i, y_i) \text{ for } i = j \\ w_1(x_i) \text{ for } (i,j) \text{ such that } \mathbf{x}_j = \mathbf{x}_i + [k, 0] \text{ for } k \in \mathbb{Z} : k \in \{x_i + 1, N_x\} \\ w_2(x_i) \text{ for } (i,j) \text{ such that } \mathbf{x}_j = \mathbf{x}_i + [-1, 0] \\ w_3(x_i, y_i) \text{ for } (i,j) \text{ such that } \mathbf{x}_j = \mathbf{x}_i + [0, 1] \\ w_4(y_i) \text{ for } (i,j) \text{ such that } \mathbf{x}_j = \mathbf{x}_i + [0, -1] \end{cases}, \tag{2}$$

We then need to solve the corresponding CME. Because the system is piece-wise constant (i.e. $\mathbf{A}$ is constant of fixed time intervals), we can pre-compute the infinitesimal generator for both the on and off state of the controller, where we define the "on" generator matrix as $\mathbf{A}_{\text{on}}$ and the "off" generator matrix as $\mathbf{A}_{\text{off}}$. Because the input is piecewise constant, given a sequence of input $u(t)$ the FSP solution can be iteratively evaluated on the constant time intervals,

$$\mathbf{p}(t + \Delta t) = \begin{cases} \exp\{\mathbf{A}_{\text{on}} \Delta t\} \mathbf{p}(t) \text{ if } u(s) = 1 \text{ for } s \in (t, t + \Delta t) \\ \exp\{\mathbf{A}_{\text{off}} \Delta t\} \mathbf{p}(t) \text{ if } u(s) = 0 \text{ for } s \in (t, t + \Delta t). \end{cases} \tag{3}$$

The matrix exponential is a notoriously expensive operation; however the product of the matrix exponential and the probability vector can be efficiently evaluated using Krylov subspace methods. We used a custom matrix exponentiation algorithm in Python based on ExpoKit [10].

#### Population Model of Protein Expression

Using standard mass-action kinetics, we find the following set of ordinary differential equations to describe the time-evolution of the mean of the population

$$\frac{dm}{dt} = k_1 - \gamma_1 \mathrm{m} \tag{4}$$

$$\frac{dx}{dt} = k_2 m - \gamma_2 x \tag{5}$$

$$\frac{dy}{dt} = k_3 u(t-\tau)x - \gamma_3 y. \tag{6}$$

In this model, $m$, $x$, and $y$ represent the concentrations of the EL222 mRNA, EL222 protein and mScarletI protein, respectively.

#### Model Calibration

Our goal was to calibrate the model such that it would be sufficient to control either the population or single-cells. We primarily used the stochastic model of the system to find model parameters which matched experimental trajectories, shown in Fig. S6. However, evaluation of the trajectory based likelihood function is very slow for these types of models, and therefore we used a combination of quantitative maximum likelihood based fitting along with hand-tuning to find the presented parameters in Fig. S6 and Tables S1 and S2.

| $k_1$ | $\gamma_1$ | $k_2$ | $\gamma_2$ | $k_3$ | $\gamma_3$ | $\tau$ |
|---|---|---|---|---|---|---|
| [molecules.min$^{-1}$] | [min$^{-1}$] | [min$^{-1}$] | [min$^{-1}$] | [min$^{-1}$] | [min$^{-1}$] | [min] |
| 0.05 | 10 | 20 | 0.0045 | 0.0125 | 0.004 | 36 |

Table S1: Deterministic model parameters

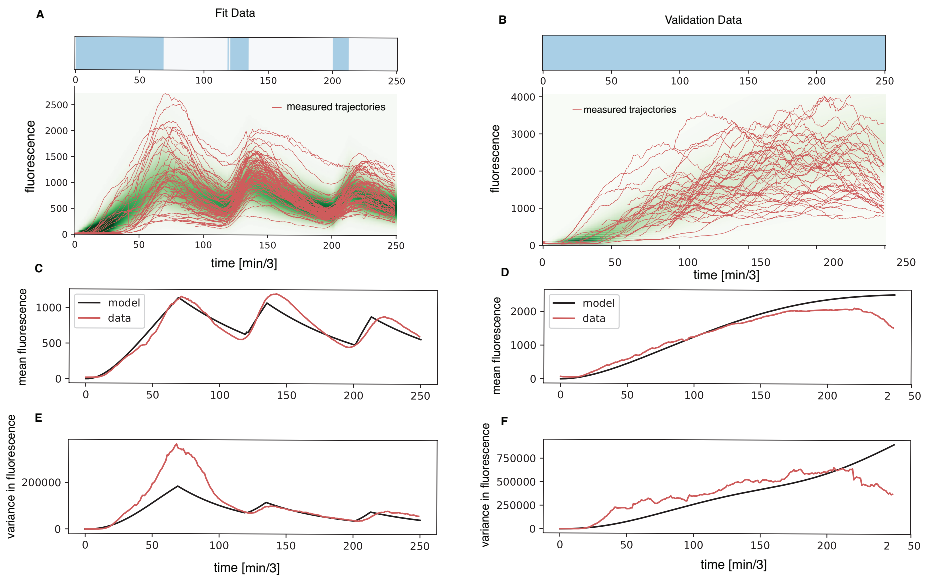


Figure S6: Model calibration and validation. (A) Probability (green heat map) and measured experimental trajectories used to calibrate the stochastic model of mScarletI production. (C-D) Corresponding mean and variance of measured data and the model. (B,D,F) Model validation on constant stimulation data. Prediction of the probability distributions (B), mean (D) and variance (F) using the parameters found by fitting the trajectories in panel A.

| $k_1$ | $\gamma_1$ | $b$ | $k_3$ | $\gamma_3$ | $\tau$ |
|---|---|---|---|---|---|
| [molecules.min$^{-1}$] | [min$^{-1}$] | [min$^{-1}$] | [min$^{-1}$] | [min$^{-1}$] | [min] |
| 0.05 | 0.0045 | 0.5 | 0.0125 | 0.004 | 36 |

Table S2: Stochastic model parameters

### Fluorescent Measurement Noise

We assume that fluorescence measurements noise is Gaussian. Therefore, if the number of fluorescent molecules in a given cell is known to be $n_i$, the fluorescence measurements are taken to be distributed as

$$f_{\mathrm{FP}}(z) = \frac{1}{\sqrt{2n_i\pi\sigma_{\mathrm{FP}}^2}} \exp\left(-\frac{(z - n_i\mu_{\mathrm{FP}})^2}{2n_i\sigma_{\mathrm{FP}}^2}\right), \tag{7}$$

where $\mu_{\mathrm{FP}}$ and $\sigma_{\mathrm{FP}}^2$ are the mean and variance of the fluorescence distribution. For this work, we calibrated the values of $\mu_{\mathrm{FP}}$ and $\sigma_{\mathrm{FP}}^2$ along with the kinetic parameters for the models, described below. Values were taken as $\mu_{\mathrm{FP}} = \sigma_{\mathrm{FP}}^2 = 50$. For the stochastic single-cell control problem, we will use Eq. 7 to estimate the number of molecules in the cell. From the population (deterministic) model standpoint, the Gaussian measurement noise fits well into the common framework of Kalman filtering, which we describe in the next section.

#### Kalman Filtering for population state estimation

For the population control of the mean, we used the deterministic model described in Eq. 4. This model describes the time evolution of the three species, $\mathbf{x} = [m, x, y]$, where $y$ is the fluorescent protein in units of number of molecules. We discretized the system in time according to the measurement times, $k\Delta t$ for $k \in (0, N_t)$. To estimate the state at each $k\Delta t$ using the measurement $\tilde{y}(k\Delta t) \equiv \tilde{y}_k$, we used a Kalman filter. The process noise $\mathbf{P}_k$ follows from the assumption of Poisson noise across the population, and is given by

$$\mathbf{P}_k = \begin{bmatrix} m(k\Delta t)/N_c & 0 & 0 \\ 0 & x(k\Delta t)/N_c & 0 \\ 0 & 0 & y(k\Delta t)/N_c \end{bmatrix}, \tag{8}$$

where $N_c$ is the number of cells in the population. This term comes from the assumption that the population mean is normally distributed with independent variances stemming from Poisson noise. Because we only measure the fluorescence, the observation matrix is

$$\mathbf{C} = \begin{bmatrix} 0 & 0 & 0 \\ 0 & 0 & 0 \\ 0 & 0 & \mu_{\mathrm{FP}}. \end{bmatrix} \tag{9}$$

At measurement times, state and process noise are updated:

$$\hat{\mathbf{x}}_{k|k} = \hat{\mathbf{x}}_{k|k-1} + \mathbf{K}_k(\tilde{y}_k - \mathbf{C}\hat{\mathbf{x}}_{k|k-1}) \tag{10}$$

$$\mathbf{P}_{k|k} = (\mathbf{I} - \mathbf{K}_k\mathbf{C})\mathbf{P}_{k|k-1} \tag{11}$$

where $\mathbf{K}_k$ is the Kalman gain,

$$\mathbf{K}_k = \mathbf{P}_{k|k-1}\mathbf{C}^{\mathrm{T}} \left(\mathbf{C}\mathbf{P}_{k|k-1}\mathbf{C}^{\mathrm{T}} + \sigma^2_{\mathrm{FP}}\right)^{-1}. \tag{12}$$

The observation variance $\sigma^2_{\mathrm{FP}}$ comes directly from Eq. 7.

#### Bayesian state estimation for single cells using the FSP

Because the Kalman filtering approach assumes that everything is Gaussian, and we know that the process is non-Gaussian, and we cannot make use of the central limit as above, we now turn to an approach for state estimation which can be applied to single cell fluorescent measurements. For simplicity, suppose we start from the estimated state of the cell, which is distributed according to $\mathbf{p}((k-1)\Delta t) = [p_0, p_1, ..., p_N]$. This distribution can be propagated from $(k-1)\Delta t$ to $k\Delta t$ according to $u(t)$ and Eq. 3. In the Bayesian sense, $\mathbf{p}(k\Delta t)$

is a prior distribution on the current state $\mathbf{x}$. Because we have a discrete state model, and we aim to estimate the number of protein $Y$, we need to marginalize over the variables which we do not observe. Recall that the stochastic model only has two variables, and therefore the marginal prior on the observation is

$$p_Y(k\Delta t) = \sum_{i\in\mathcal{X}} p_{i,j}(k\Delta t). \tag{13}$$

The likelihood of the measurement comes from Eq. 7, which gives the probability of $\tilde{y}_k$ conditioned on a given number of protein molecules in the cell, i.e. $n_i = [1, 2, 3..., N]$ molecules of protein.

$$\ell(\tilde{y}_k|n_i;\mu_{\mathrm{FP}},\sigma^2_{\mathrm{FP}}) = \frac{1}{\sqrt{2n_i\pi\sigma^2_{\mathrm{FP}}}}\exp\left(-\frac{(\tilde{y}_k - n_i\mu_{\mathrm{FP}})^2}{2n_i\sigma^2_{\mathrm{FP}}}\right). \tag{14}$$

We assume that $\mu_{\mathrm{FP}}$ and $\sigma^2_{\mathrm{FP}}$ are known. As mentioned above, Eq. 13 is a prior assumption about how many molecules of fluorescent protein a given cell is likely to have, and is conditioned on the previous estimate of the state at $(k-1)\Delta t$. Therefore, we can find the posterior estimate of the state as

$$p(n_i, k\Delta t|\tilde{y}_k(k\Delta t)) \propto \ell(\tilde{y}_k|n_i;\mu_{\mathrm{FP}},\sigma^2_{\mathrm{FP}})p_Y(n_i, k\Delta t). \tag{15}$$

The proportionality constant is not necessary to find in this case, as we can renormalize the posterior estimate of the state $p(n_i, k\Delta t|\tilde{y}_k(k\Delta t))$.

#### Receding Horizon Model Predictive Control

For both the population model and the single cell stochastic model we implemented a receding horizon model predictive control. For a given target $T$ defined at all $k\Delta t$, we defined a cost function for each method. The target has units of molecule number of fluorescent protein, which is the same units as the state $\mathbf{x}$. For the standard population control, we defined a sum of squares cost function,

$$J_{\mathrm{pop}} = \frac{1}{K}\sum_{k=0}^{K}\left(T_k - \hat{x}_k\right)^2. \tag{16}$$

For the stochastic individual cell control, we chose a different cost function which takes into account the potentially asymmetric (non-Gaussian) estimates of the state, which is the expected absolute deviation,

$$J_{\mathrm{sc}} = \frac{1}{K}\sum_{k=0}^{K}\sum_{n=0}^{N}|T_k - n|\,\tilde{p}(n, k\Delta t). \tag{17}$$

We perform receding horizon MPC, in which we aim to optimize the cost function $J$ over the finite time horizon $H$ using the current state estimate, i.e. to find the light input sequence $u(t)$ which minimizes $J$ over the finite time horizon $[k\Delta t, k\Delta t + H]$, i.e.

$$u(t) = \arg\min_{u(t)} J\left(u(t), \hat{x}\right) \tag{18}$$

For these systems, the photostimulation is taken either to be on or off between $k\Delta t$ and $(k+1)\Delta t$, therefore there are $2^{H/\Delta t}$ possible light combinations over the horizon $H$. We optimized by exhaustively evaluating Eq. 18 for each possible light sequence.

We tested this strategy *in silico* and obtained satisfying results, as shown in Fig. S7.

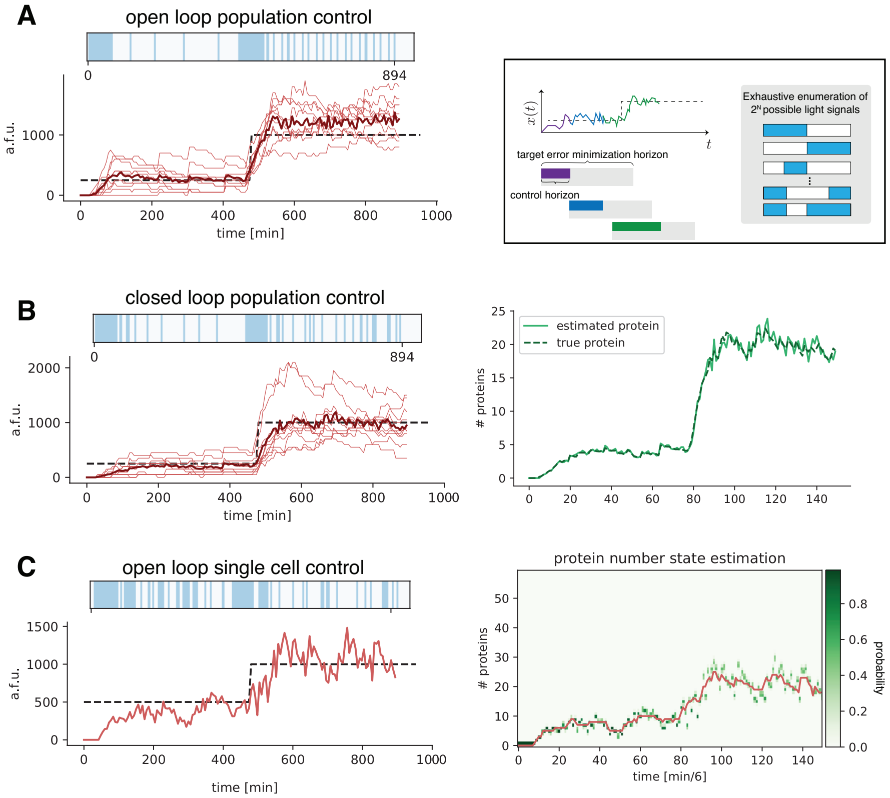


Figure S7: In silico model predictive control. (A) Open loop model predictive control of the mean particle number using the deterministic model. (B) Closed loop model predictive control and state estimation of the population mean protein number Gillespie simulations using the deterministic model. (C) Closed loop model predictive control of a single simulated trajectory using the stochastic FSP based model predictive controller.

### 5 Data Analysis

We implemented an automated pipeline to remove cells from our analysis either because of their position in the field of view or because of poor segmentation or tracking quality. Importantly, this analysis was applied only to the characterization experiments, as we only applied these cleaning strategies after the induction experiments in Fig. 2B, S4-S5 and not in real time. These cleaning steps are shown in our data analysis Jupyter notebooks. In summary, we removed cells which met one of the following criteria:

- data was collected after cells stopped growing in the microfluidic chip,
- cells were within 150 pixels of the top of the field of view, since this part of the field of view is imaged but not stimulated by the DMD, or cells were within 50 pixels of the border of the image,
- cells were present in the data set for less than 10 frames,
- cells had an area change of more than 100 square pixels between two consecutive frames.

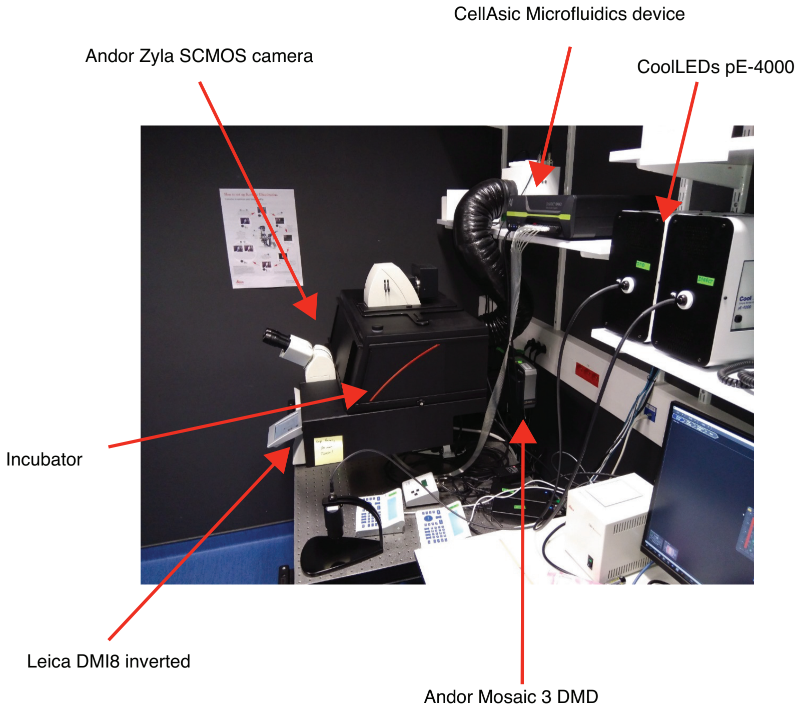


Figure S8: Microscope setup.

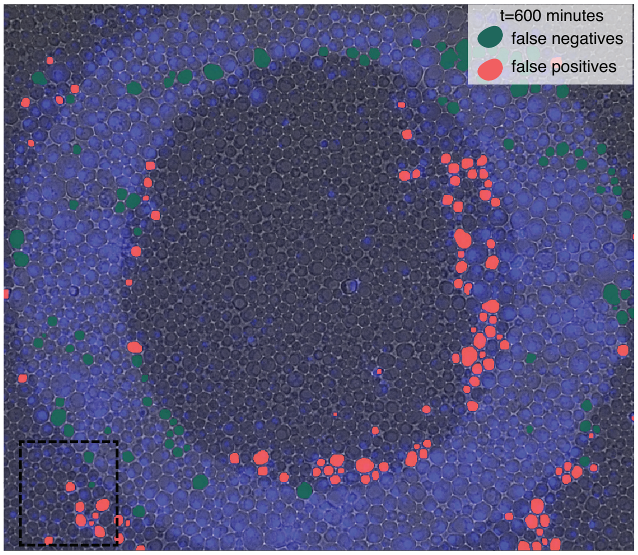


Figure S9: Recombination using geometrical patterns. The fluorescent image (blue) is superimposed with the brightfield image. Cells that are in the highest 10% of the non-targeted cells and cells that are in the lowest 10% of the targeted cells are labeled in red and green, respectively.

### References

[1] Thorsten Falk et al. U-net: deep learning for cell counting, detection, and morphometry. *Nature Methods*, 16:67–70, 2019.

[2] Jean-Baptiste Lugagne, Haonan Lin, and Mary J Dunlop. DeLTA: Automated cell segmentation, tracking, and lineage reconstruction using deep learning. *PLoS computational biology*, 16(4):e1007673, 2020.

[3] Erick Moen, Dylan Bannon, Takamasa Kudo, William Graf, Markus Covert, and David Van Valen. Deep learning for cellular image analysis. *Nature methods*, 16(12):1233–1246, 2019.

[4] François Chollet et al. Keras, 2015. https://keras.io.

[5] Martín Abadi et al. TensorFlow: Large-scale machine learning on heterogeneous systems, 2015. https://www.tensorflow.org/.

[6] Napari contributors. napari: a multi-dimensional image viewer for python, 2019.

[7] John C Crocker and David G Grier. Methods of digital video microscopy for colloidal studies. *Journal of Colloid and Interface Science*, 179(1):298–310, 1996.

[8] Cristian Versari et al. Long-term tracking of budding yeast cells in bright-field microscopy: Cellstar and the evaluation platform. *Journal of the Royal Society Interface*, 14(127), 2017.

[9] Vahid Shahrezaei and Peter S. Swain. Analytical distributions for stochastic gene expression. *Proceedings of the National Academy of Sciences*, 105(45):17256–17261, 2008.

[10] Roger B Sidje. Expokit: a software package for computing matrix exponentials. *ACM Transactions on Mathematical Software (TOMS)*, 24(1):130–156, 1998.
